# Deleterious mutations in a heterozygous chromosome result in recessive lethality in silkworm semiconsomic strains

**DOI:** 10.1101/2022.12.22.521575

**Authors:** Kenta Tomihara, Saori Tanaka, Susumu Katsuma, Toru Shimada, Jun Kobayashi, Takashi Kiuchi

**Affiliations:** Department of Agricultural and Environmental Biology, Graduate School of Agricultural and Life Sciences, The University of Tokyo, Bunkyo, Tokyo 113-8657, Japan; Department of Biological and Environmental Sciences, Graduate School of Sciences and Technology for Innovation, Yamaguchi University, Yamaguchi, Yamaguchi 753-8515, Japan; Yokohama Plant Protection Station, Ministry of Agriculture, Forestry and Fisheries, Yokohama, Kanagawa 231-0801, Japan; Department of Life Science, Faculty of Science/Graduate School of Science, Gakushuin University, Toshima, Tokyo 171-8588, Japan

**Author notes:** Corresponding author, Kenta Tomihara, Takashi Kiuchi.

**Keywords:** *Bombyx mori*, *Bombyx mandarina*, semiconsomic strain, deleterious mutation, *imaginal discs arrested* (*ida*), *TATA Box Binding Protein-associated factor 5* (*Taf5*)

## Abstract

In this study, we found two embryonic lethal mutations, *t04 lethal* (*l-t04*) and *m04 lethal* (*l-m04*), in semiconsomic strains T04 and M04, respectively. In these semiconsomic strains, the entire diploid genome, except for one chromosome 4 of the wild silkworm *Bombyx mandarina*, is substituted with chromosomes of the domesticated silkworm *B. mori*, and *l-t04* and *l-m04* mutations are located on *B. mandarina-derived* chromosome 4. The mutations responsible for the *l-t04* and *l-m04* were identified as the *Bombyx* homolog of *imaginal discs arrested (Bmida*) and *TATA Box Binding Protein-associated factor 5 (BmTaf5*), respectively. These findings indicate that both mutations were independently introduced during or after the development of semiconsomic strains. We conclude that the recessive embryonic lethality in the T04 and M04 strains is due to deleterious mutations produced in *B. mandarina*-derived chromosome 4.

**Graphical abstract:** 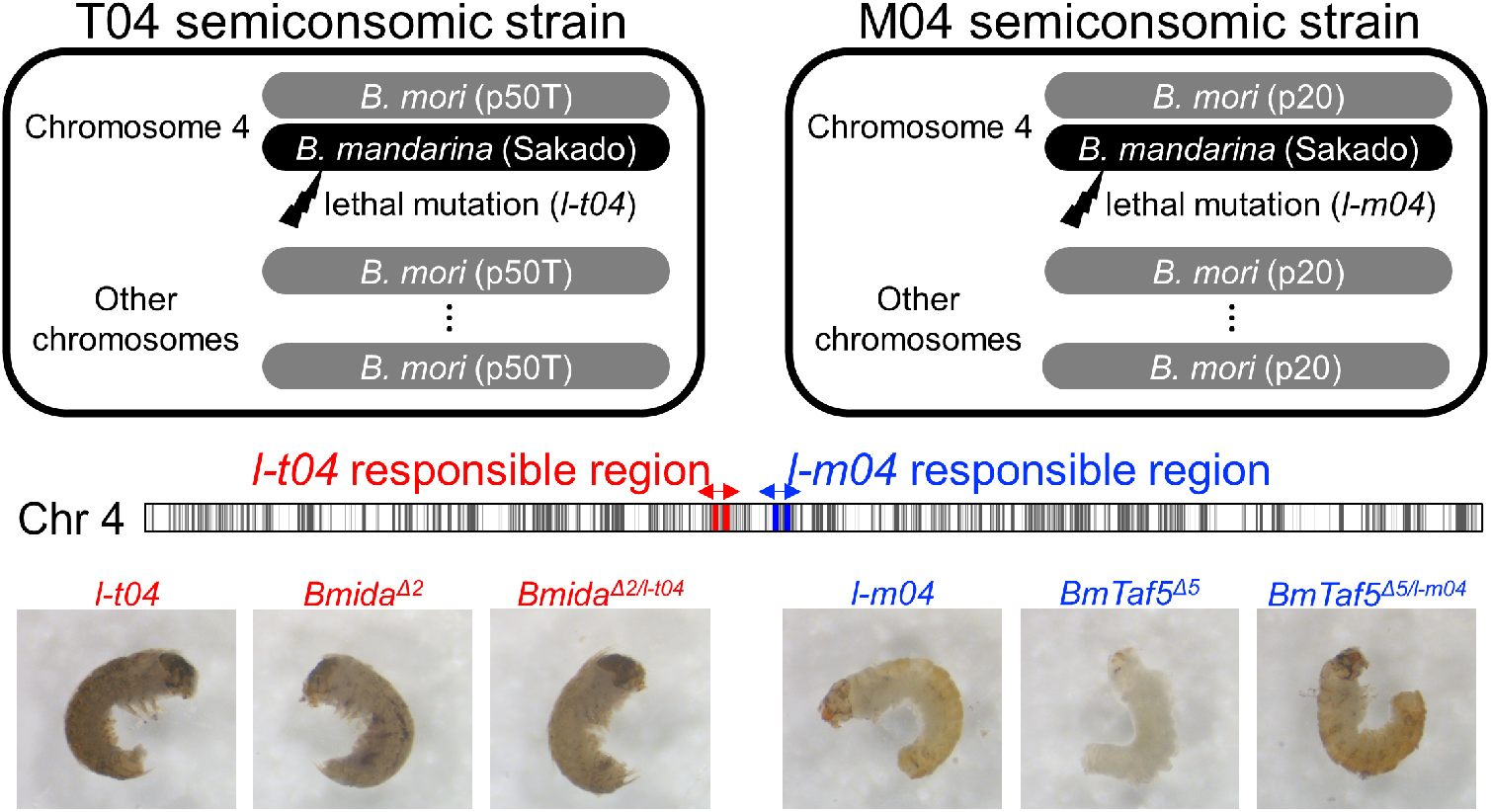

## Introduction

Mayr (1942) proposed that “Species are groups of actually or potentially interbreeding natural populations, which are reproductively isolated from other such groups.” Reproductive isolation can be classified into two broad categories: prezygotic and postzygotic isolations. Prezygotic isolation is achieved by defects before fertilization, such as differences in habitat (Estrada and Jiggins, 2002), mating time (Miyatake *et al*., 2002), behavior (Honda-Sumi, 2005), pheromone (Smadja and Butlin, 2008), and morphology (Jiggins *et al*., 2001; Kubota *et al*., 2013) among species. Postzygotic isolation is achieved by defects after fertilization and is caused by extrinsic and intrinsic reasons (Nosil, 2013). Extrinsic postzygotic isolation is mainly caused by natural selection, depending on the ecological setting. Intrinsic postzygotic isolation includes hybrid (F_1_) inviability (Sawamura *et al*., 1995; Presgraves *et al*., 2003; Brideau *et al*., 2006) and sterility (Ting *et al*., 1998). The hybrid breakdown is another type of intrinsic postzygotic isolation, which is a lethality, sterility, and weakness observed in the F2 or later generations (Stebbins, 1950). Intrinsic postzygotic isolations can be explained by negative interactions between two or more genetic loci, which is known as the Dobzhansky–Muller model (Dobzhansky, 1934; Muller, 1942).

The domesticated silkworm, *Bombyx mori*, and the wild silkworm, *B. mandarina*, are significantly close relative species. In sericulture farms, *B. mori* moths rarely encounter *B. mandarina* moths because *B. mori* pupae in cocoons are killed before emergence to obtain silk. Because *B. mori* males cannot fly and have difficulties coupling with *B. mandarina* females, only the mating combination of female *B. mori* and male *B. mandarina* is possible in natural conditions (Nakamura *et al*., 1997). Note that the copulation rate of this mating combination is still lower than in intraspecies mating (Nakamura *et al*., 1997). Furthermore, even if the eggs are oviposited in the field, the survival rate of F_1_ larvae is almost zero under natural conditions (Kômoto *et al*., 2016). It is indicated by these observations that *B. mori* and *B. mandarina* are prezygotically and postzygotically isolated for extrinsic reasons. Indeed, gene flow is not observed from *B. mori* into wild *B. mandarina* populations (Yukuhiro *et al*., 2011, 2012). Also, no F_1_ hybrid moths were collected using pheromone traps around sericulture farms (Kômoto *et al*., 2016). However, *B. mori* and *B. mandarina* can be crossed with each other under artificial conditions (Toyama, 1909; Nakamura *et al*., 1997), and their F_1_ progenies are fertile. Therefore, it has been thought that no intrinsic postzygotic isolation exists between *B. mori* and *B. mandarina*.

Consomic strains, also known as chromosome substitution strains, are inbred strains in which a chromosome pair is replaced by the corresponding chromosome pair of another strain (Fig. 1A). Such consomic strains are helpful for the comparative genomic and phenotypic studies of two strains or closely related species. Fujii *et al*. (2021) established a series of semiconsomic strains whose entire diploid genome, except for one targeted chromosome of *B. mandarina*, was replaced with *B. mori* chromosomes (Fig. 1A). T04 is one of these semiconsomic strains in which one chromosome 4 from the *B. mandarina* Sakado strain was placed on the *B. mori* p50T strain background (Fig. 1B). M04 is another semiconsomic strain in which one chromosome 4 from the *B. mandarina* Sakado strain was placed on the background of the *B. mori* p20 strain (Fig. 1B). Here we intercrossed T04 individuals with each other and found that about a quarter of F_1_ progeny eggs did not hatch. A recessive lethal gene was suggested to be located on *B. mandarina-derived* chromosome 4, which we named *t04 lethal (l-t04*) (Fig. 1B). We also found a recessive lethal gene located on *B. mandarina-derived* chromosome 4 in the M04 strain, which we named *m04-lethal* (*l-m04*) (Fig. 1B). These observations imply that a hybrid breakdown may exist between *B. mori* and *B. mandarina*.

**Figure 1:**
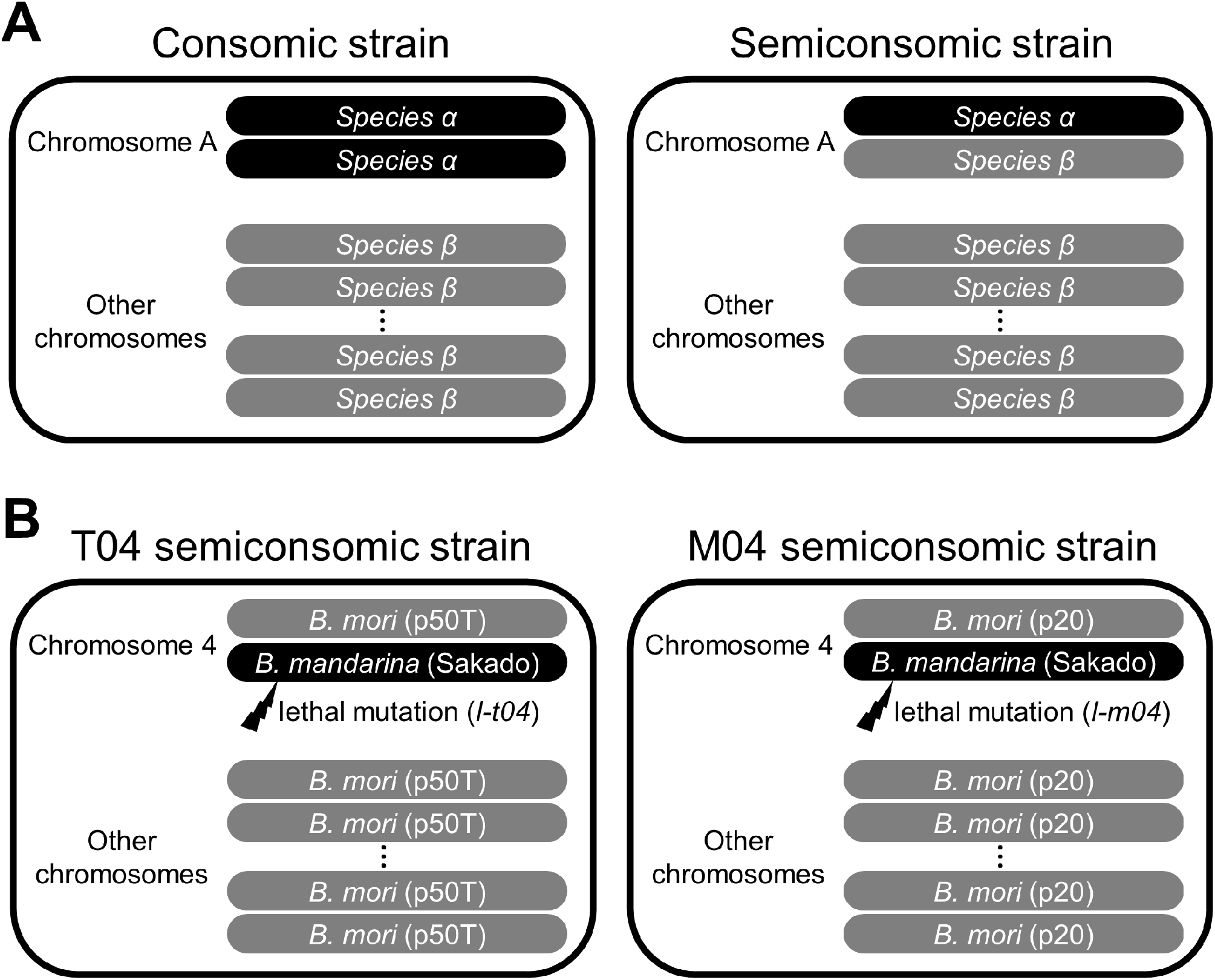
Schematic representation of consomic and semiconsimic strains. (A) Chromosomal composition of consomic and semiconsimic strains. (B) Chromosomal composition of T04 and M04 semiconsimic strains.

Alternatively, a different hypothesis can be proposed. Irreversible deleterious mutations tend to accumulate in individuals without recombination, because fewer or different combinations of mutations may be offered to offspring by recombination (i.e., Muller’s ratchet; Muller 1964). Additionally, it was shown in previous studies that deleterious mutations accumulated in a chromosome that was kept heterozygous after generations because recessive deleterious mutations did not experience selection (Mukai, 1964; Mukai *et al*., 1972). Genetic recombination does not occur in females in *B. mori* (Tanaka, 1913), and *B. mandarina-derived* chromosomes have not experienced selection during the development and maintenance of semiconsomic strains because they were transmitted to offspring through heterozygous females (Fujii *et al*., 2021). Thus, deleterious mutations might have accumulated in *B. mandarina*-derived chromosome 4 in the T04 and M04 strains, resulting in recessive lethality.

We identified the genes responsible for the *l-t04* and *l-m04* mutations to elucidate our hypotheses in this study. We showed that the lethalities in the T04 and M04 strains are not due to the hybrid breakdown between *B. mori* and *B. mandarina* but an accumulation of deleterious mutations in *B. mandarina*-derived chromosome 4. This is because the mutations responsible for *l-t04* and *l-m04* were independent and introduced during or after the development of semiconsomic strains.

## Materials and Methods

### Silkworm strains

The *B. mori* strain p50T, a single-paired descendant of individuals of strain Daizo, is maintained at our laboratory. The *B. mandarina* strain Sakado was originally collected in Sakadocity, Saitama, Japan, in 1982. Since then, it has been maintained at our laboratory by sib-mating. The semiconsomic strains T04 and M04 were provided by Kyushu University (Fukuoka, Japan) with support from the National Bioresource Project (NBRP; http://silkworm.nbrp.jp). All larvae were reared on mulberry leaves or an artificial diet (SilkmatePS [NOSAN, Japan]) at 25°C, 12L/12D condition.

### Microscope observation of early embryos

The embryos were fixed by incubating them in distilled water at 74°C for 4 min. Embryos in eggs were obtained by removing eggshells with fine tweezers under a microscope (Leica S9D; Leica Microsystems, Germany) and stained using a methylene blue solution (30% EtOH, 1% methylene blue) to allow visualization.

### Positional cloning

Polymerase chain reaction (PCR) makers showing polymorphism between *B. mori* (p50T or p20 strains) and *B. mandarina* (Sakado strain) were designed on chromosome 4, according to information on the silkworm genome database (SilkBase; Kawamoto *et al*. 2022) (Table S1). We intercrossed the T04 or M04 individuals with each other and obtained F_1_ offspring. It is difficult to distinguish genetically lethal mutant embryos and accidentally dead normal embryos from their external appearances. Therefore, only normally hatched individuals were used for positional cloning to avoid sampling errors. Genetic linkage analysis was performed using 1223 T04 × T04 F_1_ individuals and 384 M04 × M04 F_1_ individuals.

### Genomic PCR

We extracted genomic DNA from neonate larvae or adult moth legs using the HotSHOT method (Truett *et al*., 2000). Genomic PCR was performed using KOD FX Neo (TOYOBO, Japan) or KOD One (TOYOBO) following the manufacturer’s protocol.

### Isolation of total RNA and RNA sequencing (RNA-seq)

We collected T04 × T04 F_1_ eggs at 24, 72, 120, 168, and 216 hours post-oviposition (hpo) and M04 × M04 F_1_ eggs at 192 hpo. Genomic DNA and total RNA were extracted from a single embryo using TRIzol, according to the manufacturer’s protocol (Thermo Fisher Scientific, USA) (Kiuchi and Katsuma, 2022). We genotyped each embryo using primers inside the *l-t04* or *l-m04* linked regions (Table S1). Total RNA extracted from individuals whose genotypes were *l-t04/l-t04*,+^*l-04*^/+^*l-04*^, *l-m04/l-m04*, and +^*l-m04*^/+^*l-m04*^ were used for library preparation. RNA-seq libraries were sequenced using the NovaSeq 6000 platform (Ilummina, USA) with 150 bp paired-end reads. Library preparation and sequencing were performed by Novogene (China).

### RNA-seq data analysis

Low-quality reads and adapter sequences were removed using Trim Galore! (https://www.bioinformatics.babraham.ac.uk/projects/trim_galore/). We mapped the trimmed reads to the *B. mori* (p50T strain) reference genome (Kawamoto *et al*., 2019) using STAR (Dobin *et al*., 2013). The expected read count and transcripts per million (TPM) values for each gene were calculated using RSEM (Li and Dewey, 2011). Differential expression (DE) analysis was performed using the edgeR R package (Robinson *et al*., 2010).

### CRISPR/Cas9 mediated knockout (KO)

Unique target sites in the *B. mori* genome were designed using CRISPRdirect (https://crispr.dbcls.jp/; Naito *et al*. 2015). A single guide RNA (sgRNA) was synthesized using the MEGAscript T7 Transcription Kit (Thermo Fisher Scientific) with a primer set listed in Table S1, following the methods described in Bassett *et al*. (2013). In the case of sgRNA injection, a mixture of sgRNA (100 ng/μL) and Cas9 Nuclease protein NLS (300 or 600 ng/μL; NIPPON GENE, Japan) in injection buffer (100 mM KOAc, 2 mM Mg(OAc)_2_, 30 mM HEPES-KOH; pH 7.4) was prepared. In the case of CRISPR-RNA (crRNA) injection, a mixture of crRNA (200 ng/μL; FASMAC, Japan; Table S1), trans-activating crRNA (tracrRNA) (200 ng/μL; FASMAC; Table S1), and Cas9 Nuclease (300 or 600 ng/μL) in injection buffer was prepared. The mixture was injected into nondiapause p50T eggs within 6 hpo (Yamaguchi *et al*., 2011; Kiuchi and Katsuma, 2022). The injected embryos were incubated at 25°C in a humidified Petri dish until hatching.

Adult generation 0 (G_0_) moths were crossed with wild-type moths to obtain generation 1 (G_1_). Genomic DNA extraction and PCR were performed using the HotSHOT and KOD One as descrived above. Mutations at the targeted sites were detected by heteroduplex mobility assay using the MultiNA microchip electrophoresis system (SHIMADZU, Japan) (Ota *et al*. 2013; Ansai *et al*. 2014). G_1_ individuals carrying a heterozygous KO mutations were intercrossed with each other to generate homozygous KO generation 2 (G_2_) individuals. The PCR products from the targeted site in G2 were Sanger sequenced using a 3130xl Genetic Analyzer (Applied Biosystems, USA) or FASMAC sequencing service (Japan). The mutant lines are maintained as heterozygous stocks when a homozygous mutation is lethal.

### Sequencing analysis of wild *B. mandarina*

Genomic DNA of wild *B. mandarina* collected from 39 different locations in Japan was provided by Kyushu University with support from the National BioResource Project (NBRP; https://shigen.nig.ac.jp/silkwormbase/KuwakoLocMap2.do; Fig. S1). Genomic PCR and sequencing were performed as described above.

### Phylogenetic analysis

Protein sequences from humans (*Homo sapiens*), the fruit fly (*Drosophila melanogaster*), nematode (*Caenorhabditis elegans*), and yeast (*Saccharomyces cerevisiae*) were obtained from Uniprot-KB/SwissProt database. Protein sequences of *B. mori* were obtained from SilkBase. For each protein sequence dataset, BLASTP was performed to collect proteins related to the query sequence (E-value < 1e–40). All sequences were aligned using the MUSCLE program (Edgar, 2004), and a phylogenetic tree was built using MEGA X software (Kumar *et al*., 2018) and the maximum likelihood method based on the Jones–Taylor–Thornton (JTT) matrix model. Initial trees for the heuristic search were obtained automatically by applying neighbor-joining and BioNJ algorithms to a matrix of pairwise distances estimated using the JTT model. The topology was selected with a superior log likelihood value with 1000 bootstrap replicates.

## Results

### Characterization of the *l-t04* phenotype

We intercrossed T04 with each other and found that about a quarter of the F_1_ progenies were unhatched (Fig. 2A–B). Because eggs obtained from T04 females crossed to wild-type p50T males hatched normally, a recessive lethal mutation, which we named *l-t04*, was predicted to be located on *B. mandarina-derived* chromosome 4 (Fig. 1B). To investigate the lethal stage of *l-t04* mutant embryos, we removed eggshells from the dead embryos. Most of the dead embryos were at the body pigmentation stage I (Miya, 2003), corresponding to 192–216 hpo (Fig. 2C).

**Figure 2:**
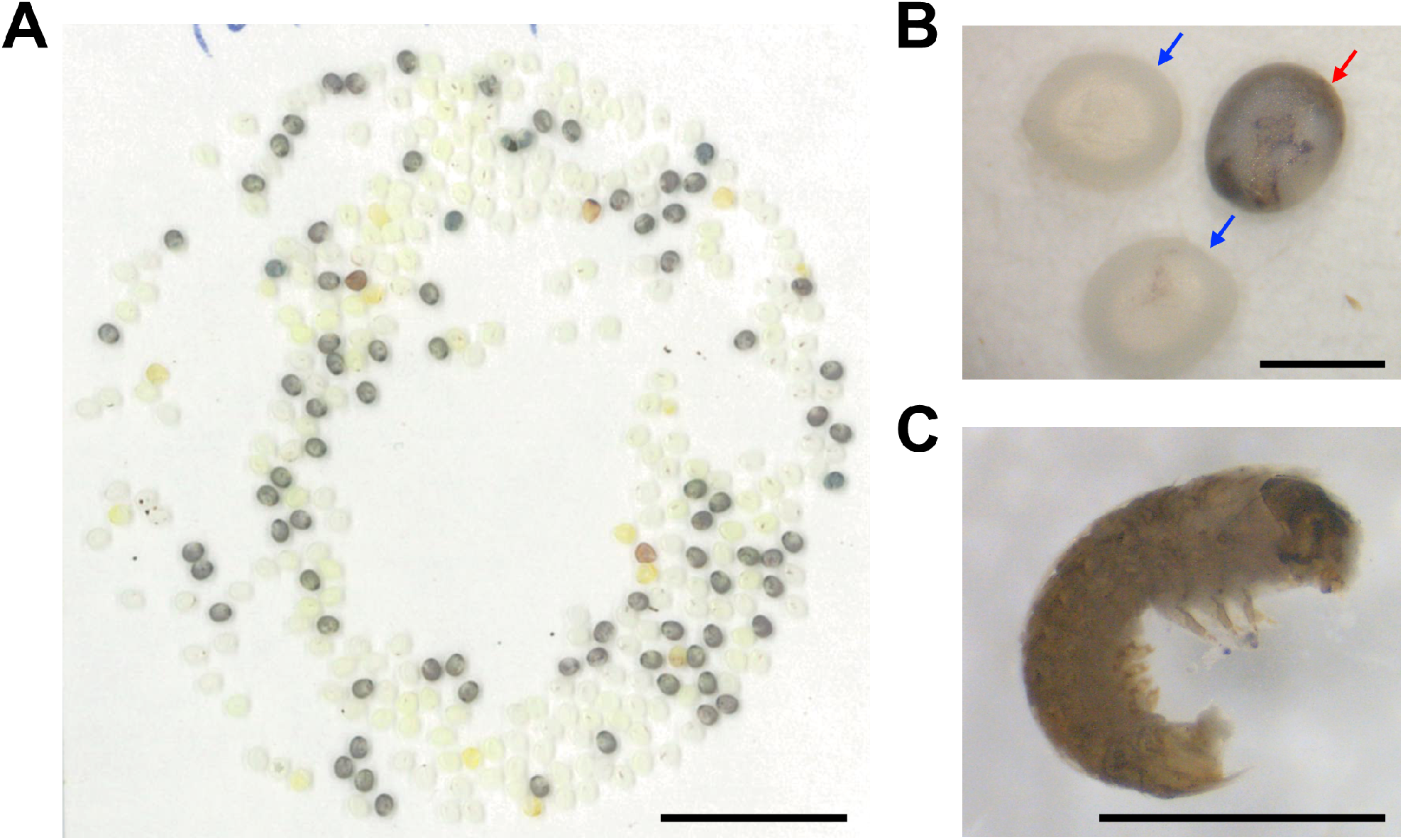
Embryonic lethality in semiconsimic strain T04. (A) Eggs laid by T04 females crossed with T04 males. (B) Enlarged view of the eggs. Blue and red arrows indicate hatched eggshells and lethal eggs, respectively. (C) Dead embryos caused by the *l-t04* mutation. Scale bars: 1 cm (A) and 1 mm (B–C).

### Positional cloning of the *l-t04* mutation

We designed PCR makers showing polymorphism between p50T and Sakado (Table S1) and performed linkage analysis using T04 × T04 F_1_ progenies. The *l-t04*-linked region was narrowed between PCR markers, Bmo04-07.97m01 and Bmo04-08.13m01 (Fig. 3A–B). This region was estimated to be approximately 153 kb, containing 13 gene models according to recent *B. mori* genome information (Table S2) (Kawamoto *et al*., 2019).

**Figure 3:**
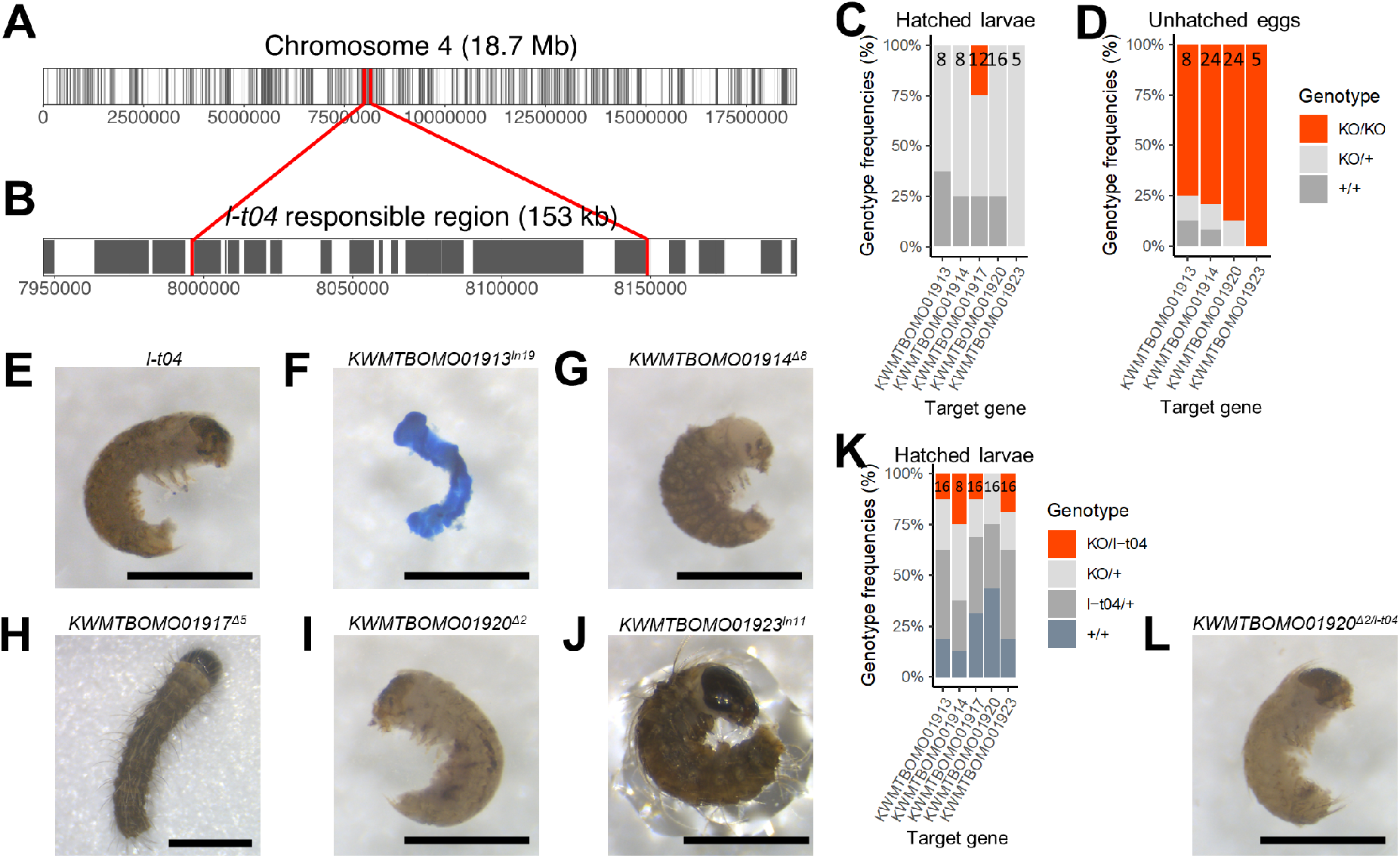
Identification of the gene responsible for the *l-t04* mutation. (A–B) Predicted gene models (gray rectangles) on chromosome 4 (A) and the *l-t04* candidate region (B). The red vertical lines indicate the borders of the region responsible for the *l-t04* locus narrowed by positional cloning. (C– D) Genotype distribution in hatched larvae (C) and unhatched eggs (D) in progenies between heterozygous KO moths. The number indicates the sample size. (E–J) Dead embryos or normally hatched larvae in *l-t04* (E), *KWMTBOMO01913^In19^* (F), *KWMTBOMO01914^Δ8^* (G), *KWMTBOMO01917^Δ5^* (H), *KWMTBOMO01920^Δ2^* (I), and *KWMTBOMO01923^In11^* (J) mutants. *KWMTBOMO01917^Δ5^* mutants hatched normally, whereas the other four KO mutants died at the embryonic stage. *KWMTBOMO01913^In19^* embryo was stained using methylene blue to allow visualization. (K) Genotype distribution in hatched larvae obtained from KO heterozygous individuals crossed to T04 individuals. (L) Dead F_1_ embryos were obtained from complementary crosses between *KWMTBOMO01920^Δ2/+^* females and T04 males. The genotypes of each embryo and larva were confirmed by genomic PCR using the primers in Table S1. Scale bars: 1 mm.

### Screening of the responsible gene for the *l-t04* mutation by CRISPR/Cas9-mediated KO

We designed unique guide RNA target sites in five of the 13 candidate genes (*KWMTBOMO01913, KWMTBOMO01914, KWMTBOMO01917, KWMTBOMO01920*, and *KWMTBOMO01923;* Fig. S2, Table S1). We successfully obtained KO individuals with homozygous mutations for each target gene (Methods, Fig. S2). KO mutants *KWMTBOMO01913^In19^, KWMTBOMO01914^Δ8^, KWMTBOMO01917^Δ5^, KWMTBOMO01920^Δ2^*, and *KWMTBOMO01923^In11^* were used in further studies. These mutants have mutations that result in frame shifts and premature stop codons.

The *KWMTBOMO01917^Δ5^* mutant was viable (Fig. 3C), and the visible phenotype was comparable to that of the wild type throughout its life stage (Fig. 3H, Fig. S3). On the other hand, *KWMTBOMO01913^In19^, KWMTBOMO01914^Δ8^, KWMTBOMO01920^Δ2^*, and *KWMTBOMO01923^In11^* mutants were lethal at the embryonic stage (Fig. 3C–D). Therefore, we removed eggshells from the dead embryos and investigated their lethal stages (Fig. 3F–G and I–J). *KWMTBOMO01913^In19^* embryos died at the early developmental stage (Fig. 3F), whereas *KWMTBOMO01923^In11^* embryos died just before hatching (Fig. 3J). *KWMTBOMO01914^Δ8^* embryos exhibited malformation: the size of their heads was normal, but their bodies were severely shrunken (Fig. 3G). Furthermore, *KWMTBOMO01920^Δ2^* embryos were lethal at body pigmentation stage I, which was similar to the *l-t04* mutants (Fig. 3I).

Next, we performed a genetic complementation test by crossing T04 (*l-t04*/+) individuals with each heterozygous mutant and examined whether their progenies were lethal at the embryonic stage. *KWMTBOMO01913^In19/l-t04^, KWMTBOMO01914^Δ8/l-t04^*, and *KWMTBOMO01923^In11/l-t04^* larvae hatched normally (Fig. 3K). On the other hand, *KWMTBOMO01920^Δ2/l-t04^* individuals were lethal at body pigmentation stage I as *l-t04* homozygous mutant embryos (Fig. 3K–L). It is indicated by these results that the mutation in *KWMTBOMO01920* is responsible for the *l-t04* lethal phenotype. We termed this gene *Bombyx* homolog of *ida (Bmida*) because *KWMTBOMO01920* is homologous to *imaginal discs arrested (ida*) of *D. melanogaster*.

### The *l-t04* mutation is specific to the T04 strain

The hatchability of T04 (*l-t04/+*) × F_1_ (p50T × Sakado) progeny was normal (93.8%), suggesting that the *l-t04* mutation is not available in the *B. mandarina* Sakado strain. We then performed RNA-seq analysis using eggs collected at 24, 72, 120, 168, and 216 hpo. The expression of *Bmida* was significantly lower in *l-t04* mutants than in the wild type at 72 and 120 hpo (false discovery rate [FDR] < 0.01; Table S3, Fig. S4). We viewed the mapped RNA-seq reads on the *Bmida* gene and noticed that the *l-t04* mutant had a nonsense mutation (G1969T), which did not exist in the Sakado strain (Fig. 4, Fig. S5). We sequenced the genomic region containing the G1969T site in wild *B. mandarina* collected from 39 distinct locations in Japan (Fig. S1), but none had this mutation (Fig. S5). It is indicated by these results that the G1969T mutation in *Bmida* is specific to the T04 strain.

**Figure 4:**
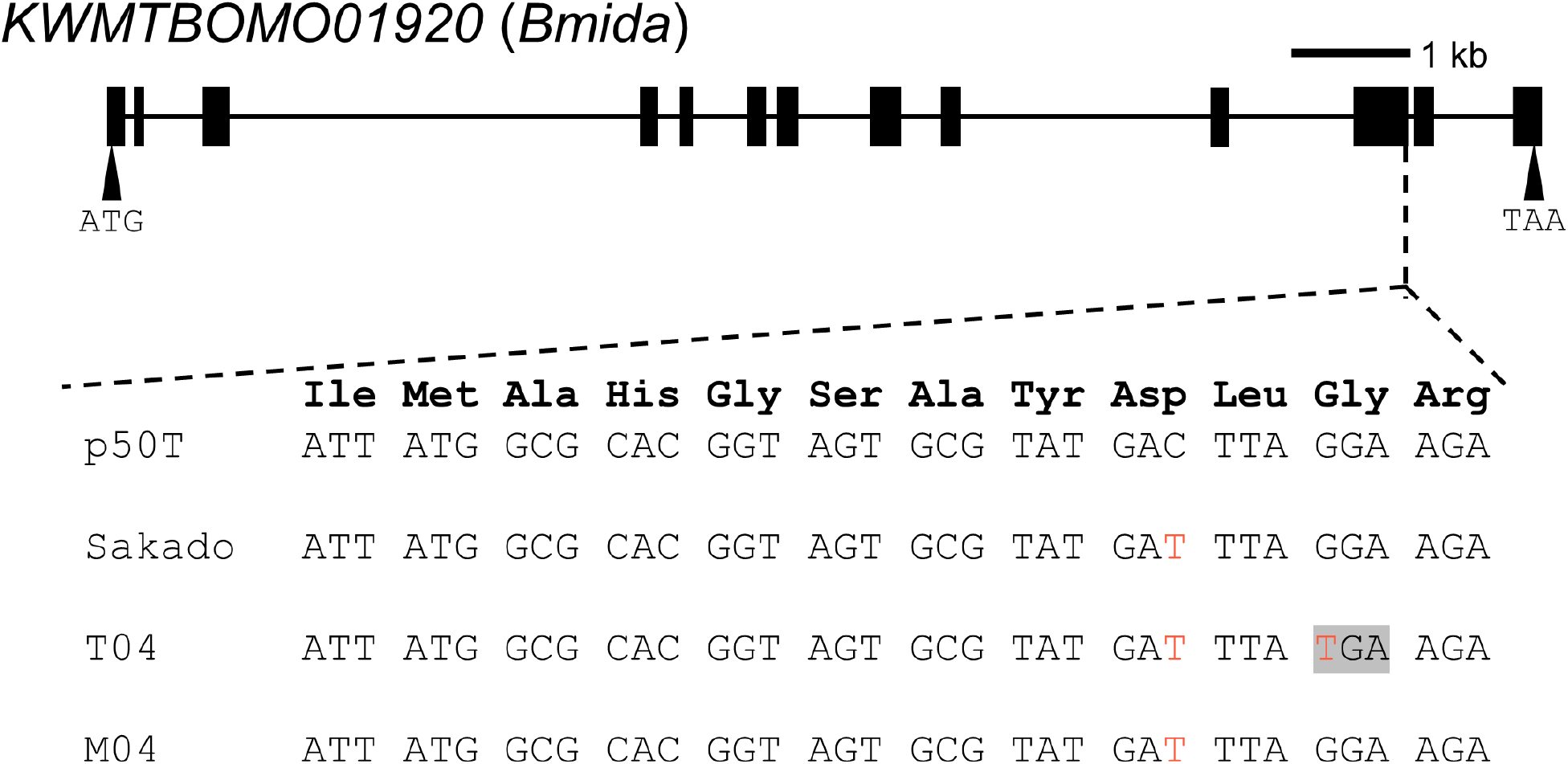
The exon structure of *KWMTBOMO01920* and alignment of *KWMTBOMO01920* gene sequences surrounding the guide RNA target site from wild-type *B. mori* strain p50T, *B. mandarina* strain Sakado, semiconsomic strains T04 (*B. mandarina-derived* allele), and M04 (*B. mandarina-derived* allele). Single nucleotide polymorphism (SNP) variants different from p50T are shown in red letters. A premature stop codon is highlighted with gray shadings.

### Characterization of the *l-m04* phenotype

About a quarter of the M04 × M04 progenies were unhatched (Fig. 5A–B). The unhatched embryos died at head pigmentation stage I (Miya, 2003), slightly before the *l-t04* mutant lethal stage (Fig. 5C). We crossed M04 to T04 individuals and found that most of their F_1_ progenies, including individuals with homozygous *B. mandarina* at chromosome 4, hatched normally (total hatching rate 92%; Fig. S6). Additionally, the G1969T mutation was not observed in the *Bmida* of the M04 strain (Fig. 4, Fig. S5). It is suggested by these results that the lethal mutations in M04, which we named *l-m04*, and *l-t04* are independent.

**Figure 5:**
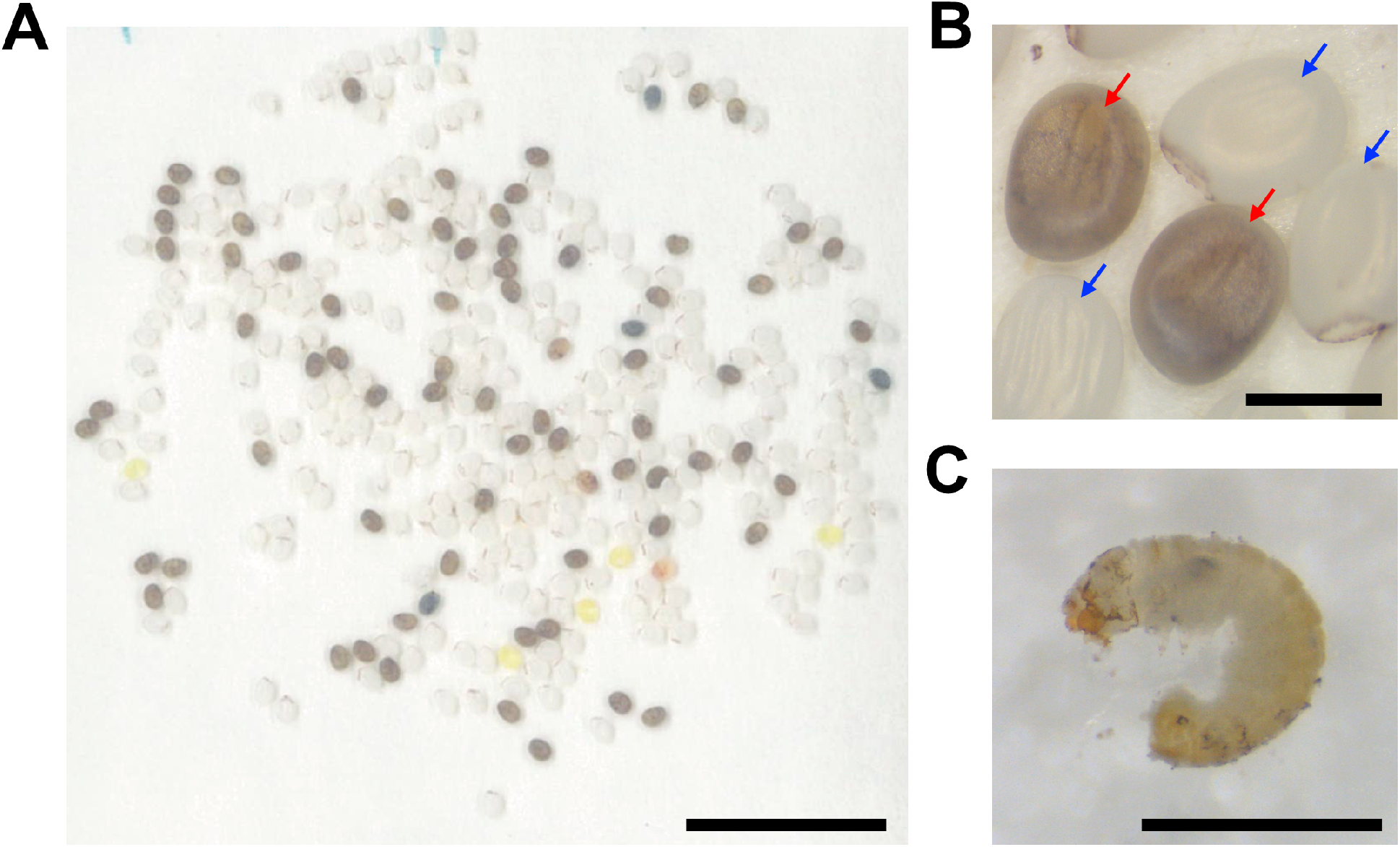
Embryonic lethality in semiconsimic strain M04. (A) Eggs laid by M04 females crossed to M04 males. (B) Enlarged view of the eggs. Blue and red arrows indicate hatched eggshells and unhatched eggs, respectively. (C) Dead embryos caused by the *l-m04* mutation. Scale bars: 1 cm (A) and 1 mm (B–C).

### Positional cloning of the *l-m04* mutation and identification of the candidate genes using RNA-seq

Using primer sets listed in Table S1, we performed a linkage analysis using M04 × M04 progenies. The *l-m04*-linked region was narrowed to between PCR markers, Bmo04-09m01 and Bmo04m-09.01m01. This region was approximately 169 kb, containing 15 gene models (Kawamoto *et al*., 2019) and about 759 kb away from the *Bmida* locus (Fig. 6A–B).

**Figure 6:**
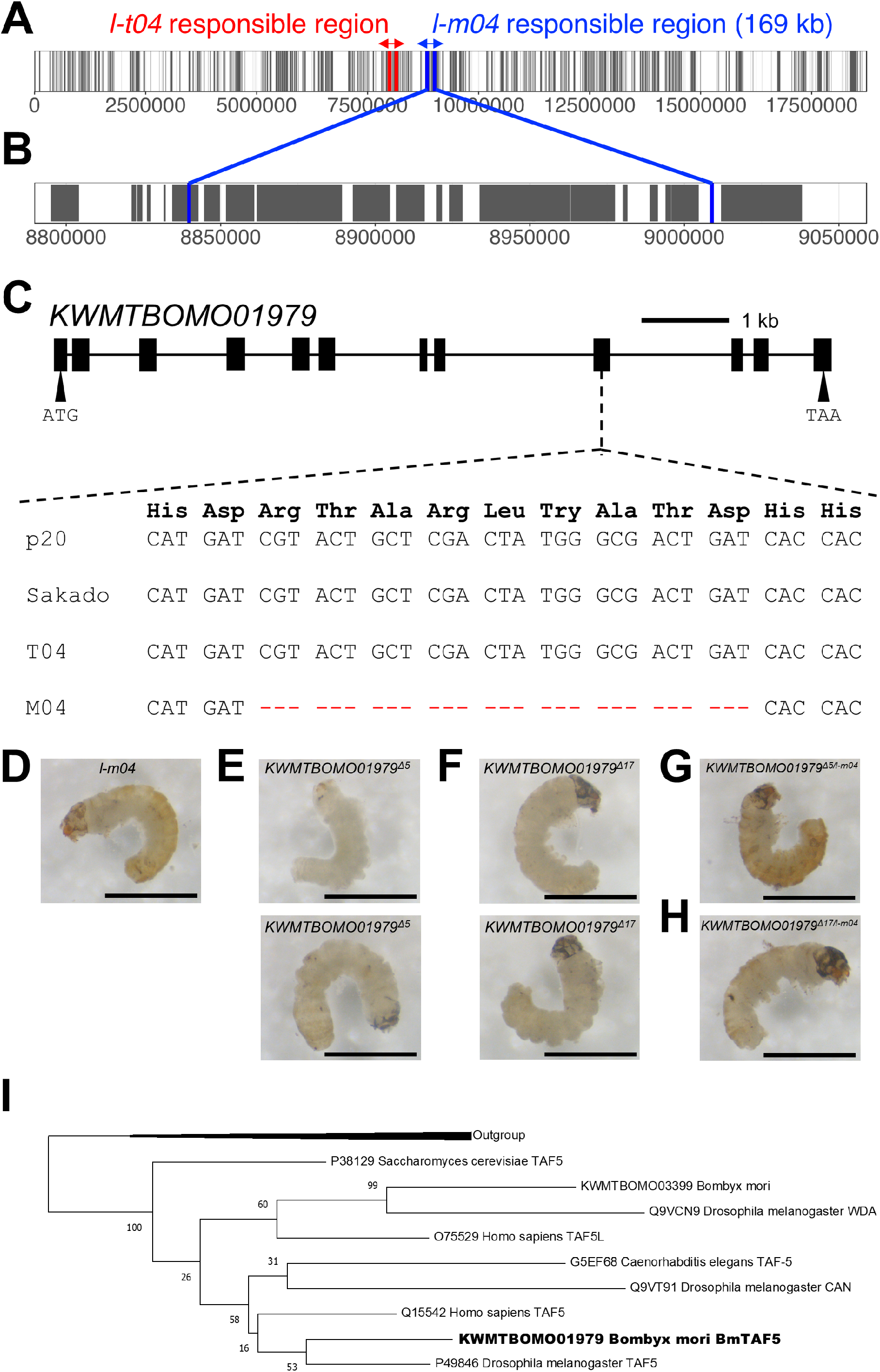
Identification of the gene responsible for the *l-m04* mutation. (A–B) Predicted gene models (gray rectangles) on chromosome 4 (A) and the *l-m04* candidate region (B). The red and blue vertical lines indicate the borders of the responsible region for the *l-t04* and *l-m04* loci, respectively. (C) The exon structure of *KWMTBOMO01979* and alignment of *KWMTBOMO01979* gene sequences surrounding the guide RNA target site from wild-type *B. mori* strain p20, *B. mandarina* strain Sakado, semiconsomic strains T04 (*B. mandarina-derived* allele), and M04 (*B. mandarina-derived* allele). SNP variants different from p20 are shown in red letters. (D–E) Dead embryos of *l-m04* (D), *KWMTBOMO01979^Δ5^* (E), and *KWMTBOMO01979^Δ17^* (F) mutants. (G–H) Dead F_1_ embryos obtained from complementary crosses between *KWMTBOMO01979^Δ5^* (G) or *KWMTBOMO01979^Δ17^* (H) females and M04 males. The genotype of each embryo was confirmed by genomic PCR using the primers shown in Table S1. (I) Phylogenetic tree of TAF5-related proteins in *H. sapience, D. melanogaster*, *C. elegans, S. cerevisiae*, and *B. mori*. Accession numbers are given with the names of the species. BmTAF5 is shown in bold letters.

Next, we performed RNA-seq analysis using total RNA extracted from embryos at 192 hpo. Although none of the genes in the responsible region were differently expressed, after checking the mapped reads, we noticed the existence of a 27 bp deletion in exon 9 of *KWMTBOMO01979* (Fig. 6C). This 27 bp deletion in *KWMTBOMO01979* was not found in parental strains p20 and Sakado or wild *B. mandarina* collected from 39 distinct locations in Japan (Fig. 6C, Fig. S7).

### Functional analysis of the *l-m04* candidate gene by CRISPR/Cas9-mediated KO

We designed a unique crRNA target site in *KWMTBOMO01979* and successfully obtained KO individuals that had homozygous mutations (*KWMTBOMO01979^Δ5^, KWMTBOMO01979^Δ9^*, and *KWMTBOMO01979^Δ17^* in Fig. S8). *KWMTBOMO01979^Δ5^* and *KWMTBOMO01979^Δ17^* were used in further studies because these mutations resulted in a frame shift and premature stop codon. *KWMTBOMO01979^Δ17^* and *KWMTBOMO01979^Δ5^* mutant embryos were embryonic lethal, but their lethal stages were not uniform and differed from the *l-m04* mutants (Fig. 6D–F, Fig. S9). We crossed *KWMTBOMO01979* heterozygous KO mutants with M04 (*l-m04/+*) and found that both *KWMTBOMO01979^Δ/l-m04^* and *KWMTBOMO01979^Δ17/l-m04^* progenies were embryonic lethal (Fig. 6G–H, Fig. S9), indicating that mutations in *KWMTBOMO01979* were responsible for the lethal phenotype in the *l-m04* mutant. Since *KWMTBOMO01979* was homologous to *D. melanogaster TATA Box Binding Protein (TBP)-associated factor 5 (Taf5*) we named this gene *BmTaf5* (Fig. 6I). The deletion of nine amino acids in the *l-m04* mutant was located at the WD-40 repeat-like domain (Fig. S10A), which was essential for protein-protein interaction (Schapira *et al*., 2017). Three of the nine amino acid residuals were conserved in the TAF5 homologs of *H. sapience, D. melanogaster*, *C. elegans*, and *S. cerevisiae* (Fig. S10B).

## Discussion

### The *l-t04* and *l-m04* mutations were independently introduced to the semiconsomic strains

In this study, we found two embryonic lethal mutations, *l-t04* and *l-m04*, in different semiconsomic strains, T04 and M04, respectively. It was suggested by the genetic complementation test between *l-t04* and *l-m04* that these two mutations were independent (Fig. S6), and their responsible genes were different (Fig. 3, Fig. 6). The lethal mutations responsible for *l-t04* and *l-m04* were not found in parental strains (p50T, p20, and Sakado) or wild *B. mandarina* collected from 39 distinct locations in Japan (Fig. 4, Fig. 6, Fig. S5, Fig. S7). This suggests that these mutations were introduced during or after the establishment of the semiconsomic strains. Thus, we concluded that the embryonic lethality observed in the T04 and M04 strains was not due to the hybrid breakdown between *B. mori* and *B. mandarina*.

Irreversible deleterious mutations tend to accumulate without recombination (Muller, 1964). Additionally, deleterious mutations accumulate in a chromosome that was kept heterozygous throughout generations because recessive deleterious mutations do not experience selection. Crossing over is absent in males in *D. melanogaster* (Morgan, 1912, 1914) and females in *B. mori* (Tanaka, 1913). Take *D. melanogaster*, for example; deleterious mutations accumulate in a chromosome that was transmitted through heterozygous males after generations (Mukai, 1964; Mukai *et al*., 1972). In establishing semiconsomic strains, *B. mandarina* females were first crossed with *B. mori* males. Subsequently, the female progenies were backcrossed to *B. mori* in every generation until all chromosomes except one were replaced by *B. mori*-derived chromosomes. Once semiconsomic strains were established, they were maintained by backcrossing the semiconsomic females to *B. mori* males (Fujii *et al*., 2021). Thus, during the establishment and maintenance of semiconsomic strains, *B. mandarina*-derived chromosomes were transmitted through heterozygous females without recombination. Also, deleterious mutations are likely to accumulate by genetic drift in a small population (i.e., mutational meltdown; Lynch and Gabriel, 1990). Indeed, during the establishment process, each semiconsomic strain was selected from a single mate cross in every generation (Fujii *et al*., 2021). Overall, our results alert researchers to the potential dangers of using laboratory animals with a heterozygous chromosome: unexpected mutations may accumulate on a heterozygous chromosome that may affect experimental outcomes. Further studies are required to investigate whether deleterious mutations accumulate in other semiconsomic strains.

### A mutation in *Bmida* is responsible for *l-t04* embryonic lethality

In this study, we identified the gene responsible for *l-t04, Bmida* (Fig. 3, Fig. 4). According to the homology to its orthologs in *D. melanogaster (ida*) and humans (*anaphase-promoting complex subunit 5; ANAPC5*), the gene responsible for *l-t04, Bmida*, is predicted to encode a subunit of the anaphase-promoting complex/cyclosome (APC/C). It is not surprising that loss-of-function mutants of *Bmida* are lethal at the embryonic stage because APC/C is essential for fundamental life activities, i.e., cyclin degradation and mitotic progression in the cell cycle (McLean *et al*., 2011). We found that the mRNA expression of *Bmida* was significantly lower in *l-t04* mutants than in the wild type at 72 and 120 hpo (Table S3), probably after maternal *Bmida* mRNA was depleted. The number of differentially expressed genes increased after 168 and 216 hpo, suggesting that the effect of *Bmida* depletion was detrimental at this time (Fig. S4). These observations consist of the lethal stage observed in the *l-t04* and *Bmida* KO mutants (192-216 hpo). On the other hand, *ida* mutants in *D. melanogaster* die during metamorphosis, in which the lethal stage is noticeably later than in *B. mori* (Bentley *et al*., 2002). This suggests that the essentialities of *Drosophila* IDA and BmIDA are slightly different. Further studies are required to reveal the functional conservation and divergence of APC/C components in insects.

### A mutation in *BmTaf5* is responsible for *l-m04* embryonic lethality

We found that nine amino acid deletions at the WD-40 repeat-like domain of *BmTaf5* were responsible for the *l-m04* lethal phenotype (Fig. 6, Fig. S10). In the budding yeast *S. cerevisiae*, TAF5 is a component of the transcription factor IID (TFIID), basal transcription factor complex, and Spt/Ada/Gcn5 acetyltransferase (SAGA) complex. TFIID is required for transcriptional initiation by RNA polymerase II (Papai *et al*., 2011), whereas SAGA functions in histone modification (Koutelou *et al*., 2010). In *S*. *cerevisiae* and the fission yeast *Schizosaccharomyces pombe*, WD-40 repeats are essential for TAF5 incorporation into TFIID and SAGA complexes (Durso *et al*., 2001; Mitsuzawa and Ishihama, 2002). On the other hand, *Taf5* homolog genes are often duplicated in other species (Fig. 6I). For example, humans have two *S. cerevisiae* TAF5 paralogs, TAF5 and TAF5 Like (TAF5L), which are incorporated into the TFIID and SAGA complexes, respectively (Helmlinger and Tora, 2017). *D. melanogaster* has three *S. cerevisiae Taf5* paralogs, *Taf5*, *cannonball* (*can*), and *will decrease acetylation (wda*). In *D. melanogaster*, TAF5 and WDA are incorporated into TFIID and SAGA complexes, respectively (Guelman *et al*., 2006), while CAN has a testis-specific function (Hiller *et al*., 2001). *Taf5* knockdown and *wda* loss-of-function mutants result in lethality at the pupal and second instar larval stages, respectively (Guelman *et al*., 2006; Neely *et al*., 2010), whereas *can* mutants are viable (Hiller *et al*., 2001). *B. mori* has two TAF5 homologs, BmTAF5 (KWMTBOMO01979) and KWMTBOMO03399, which are orthologous to *D. melanogaster* TAF5 and WDA, respectively, and lack a CAN orthologue (Fig. 6I). Overall, it is indicated by our results that nine amino acid deletions at the BmTAF5 WD-40 repeat found in the *l-m04* mutant are critical for *B. mori* embryonic development. This is probably because the function of the TFIID complex is disrupted by this mutation.

### Other essential genes for embryonic development found in this study

In *B. mori*, KO of *KWMTBOMO01913* resulted in lethality in the early embryonic stage (Fig. 3F). *KWMTBOMO01913* is homologous to *D. melanogaster c12.1* and human *CWC15* (Table S2), both of which encode a component of the spliceosome and are involved in mRNA splicing (Herold *et al*., 2009). *c12.1* KO mutants in *D. melanogaster* are lethal at the late pupal stage (Schnorrer *et al*., 2010), indicating the importance of *c12.1* in normal development. Loss-of-function mutations in *CWC15* result in embryonic lethality in the Jersey cattle *Bos taurus* (Sonstegard *et al*., 2013) and the thale cress *Arabidopsis thaliana* (Slane *et al*., 2020), suggesting that the essentiality of *CWC15* homologs is conserved in a wide range of eukaryotes.

In *B. mori*, *KWMTBOMO01914* KO mutants exhibited malformed body shrinkage and were lethal at the embryonic stage (Fig. 3G). *KWMTBOMO01914* homologs in *D. melanogaster (caper*) and humans (*RNA Binding Motif Protein 39; RBM39*) encode an RNA-binding protein that mediates alternative splicing (Table S2). In *D. melanogaster*, although semi-lethality is caused by *caper* knockdown via *ActinGal4*, the hypomorphic mutants of *caper* are viable (Olesnicky *et al*., 2017; Titus *et al*., 2021). This suggests that the essentiality of *caper* differs between *B. mori* and *D. melanogaster*. Further studies using loss-of-function mutants are required in *D. melanogaster*.

In *B. mori, KWMTBOMO01923* KO resulted in lethality at the late embryonic stage (Fig. 3J). The KWMTBOMO01923 homolog in *D. melanogaster*, Serpentine (Serp), is a chitin deacetylase and has a function in cuticle formation, such as tracheal tube development and wing differentiation (Luschnig *et al*., 2006; Wang *et al*., 2006; Zhang *et al*., 2019). *D. melanogaster Serp* mutants exhibit a tracheal tube-overextension phenotype and are embryonic lethal (Wang *et al*., 2006). This indicates the importance of this gene in normal development in insects. KWMTBOMO01923 homologs are widely found in metazoa, bacteria, and fungi (data not shown, BLASTP against NCBI non-redundant database). Intriguingly, KWMTBOMO01923 homologs are conserved in amphioxus but not in vertebrates, except for the sea lamprey *Petromyzon marinus* (data not shown, BLASTP against NCBI non-redundant database). Amphioxus has two types of chitin synthase (CS) genes (type-I and II), whereas fish and amphibians lost type-II CSs, and birds and mammals lost both types of CSs (Shi *et al*., 2020). Birds and mammals do not produce chitin, and although fish and amphibians produce chitin endogenously (Tang *et al*., 2015), it is not clear whether chitin is essential for their survival. In most vertebrates, KWMTBOMO01923 homologs have been lost, probably because chitin is not as important as in other species.

## Acknowledgement

We are grateful to Peter Andolfatto and Andrew Taverner for helpful discussions and assistance with the RNA-seq analysis. We are also grateful to Yutaka Banno and Tsuguru Fujii for providing information of semiconsomic strains. We also thank the Institute for Sustainable Agroecosystem Services, The University of Tokyo, for facilitating the mulberry cultivation and the Biotron Facility at the University of Tokyo for rearing the silkworms. This work was supported by JSPS KAKENHI grant number JP20J22954 to KT and JP18H03949 to TS.

## Data availability

RNA-seq data was submitted to DDBJ under accession numbers DRA015362 and DRA015363.

## Author contributions

KT, TS and TK designed the study. KT conducted most of the experiments and analyses. ST and JK found embryonic lethal phenotype in T04 strain and performed a pilot study. KT wrote the draft manuscript and TK revised it with the inputs of SK and TS. All authors approved the final version of the manuscript and agree to be accountable for all aspects of the work.

