## Supplemental figures for "Deleterious mutations in a heterozygous chromosome result in recessive lethality in silkworm semiconsomic strains"

*Corresponding author

A

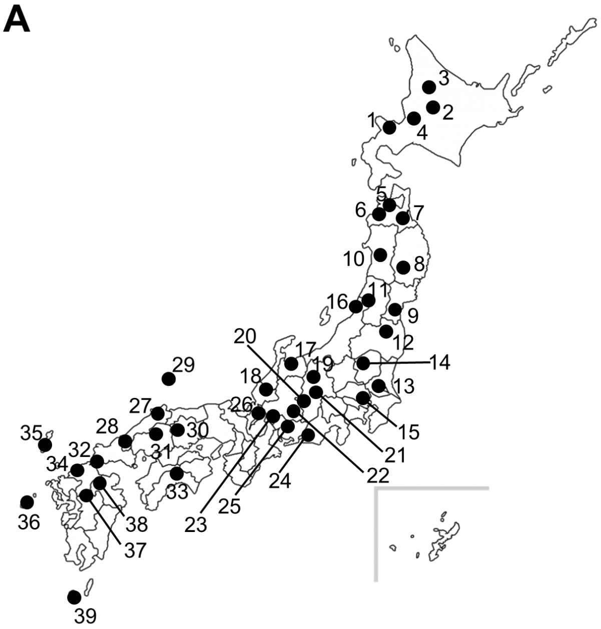


B

| Sample ID | Prefecture | Municipalities |
|---|---|---|
| 1 | Hokkaido | Sapporo |
| 2 | | Asahikawa |
| 3 | | Sibetsu |
| 4 | | Sunagawa |
| 5 | Aomori | Aomori |
| 6 | | Owani |
| 7 | | Gonohe |
| 8 | Iwate | Kitakami |
| 9 | Miyagi | Natori |
| 10 | Akita | Ogatsu |
| 11 | Yamagata | Oguni |
| 12 | Fukushima | Nihonmatsu |
| 13 | Ibaraki | Tsukuba |
| 14 | Tochigi | Nikko |
| 15 | Saitama | Saitama |
| 16 | Niigata | Itoigawa |
| 17 | Toyama | Toyama |
| 18 | Fukui | Katsuyama |
| 19 | Nagano | Azumino |
| 20 | | Kiso |
| 21 | | Hakuba |
| 22 | Gifu | Ena |
| 23 | | Kaizu |
| 24 | Shizuoka | Kosai |
| 25 | Aichi | Toyota |
| 26 | Shiga | Nagahama |
| 27 | Shimane | Unnan |
| 28 | | Tsuwano |
| 29 | | Okinoshima |
| 30 | Okayama | Niimi |
| 31 | Hiroshima | Miyoshi |
| 32 | Yamaguchi | Shimonoseki |
| 33 | Kochi | Otoyo |
| 34 | Fukuoka | Fukuoka |
| 35 | Nagasaki | Tsushima |
| 36 | | Goto |
| 37 | Kumamoto | Tamana |
| 38 | Ooita | Hita |
| 39 | Kagoshima | Yakushima |

**Figure S1**: Map (A) and table (B) of the locations where wild *B. mandarina* samples were collected.

# A

*KWMTBOMO01913*

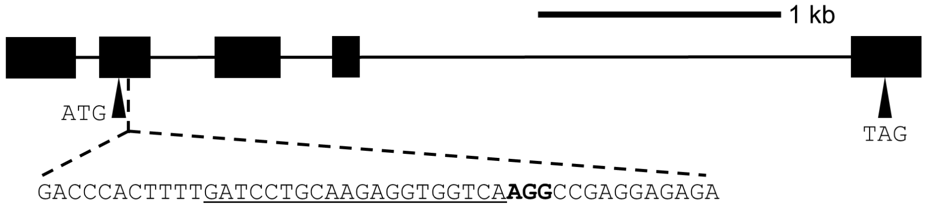


```
        Phe Asp Pro Ala Arg Gly Gly Gln Gly Arg Gly Glu Lys Asp Leu
p50T    TTT GAT CCT GCA AGA GGT GGT CAA GGC CGA GGA GAG AAA GAC TTA

Δ2      TTT GAT CCT GCA AGA GGT --T CAA GGC CGA GGA GAG AAA GAC TTA

In19    TTT GAT CCT GCA AGA GGC GGT CAA GGC CGA GGA GAG AAA GAC TTA
                            CGAGGCCGGGCCAAGGCCG
```

# B

*KWMTBOMO01914*

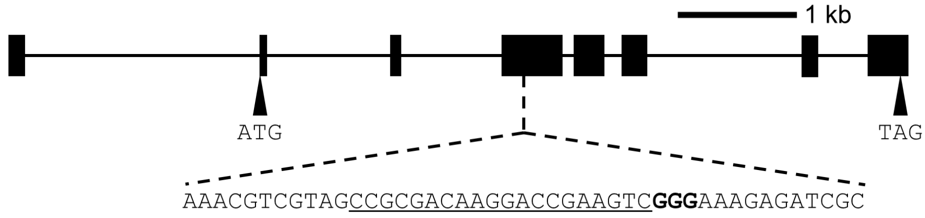


```
        Arg Ser Arg Asp Lys Asp Arg Ser Arg Glu Arg Asp Arg Ser Arg
p50T    CGT AGC CGC GAC AAG GAC CGA AGT CGG GAA AGA GAT CGC AGT CGC

Δ8      CGT AGC CGC GAC AAG GAC CGA A-- --- --- AGA GAT CGC AGT CGC
```

# C

*KWMTBOMO01917*

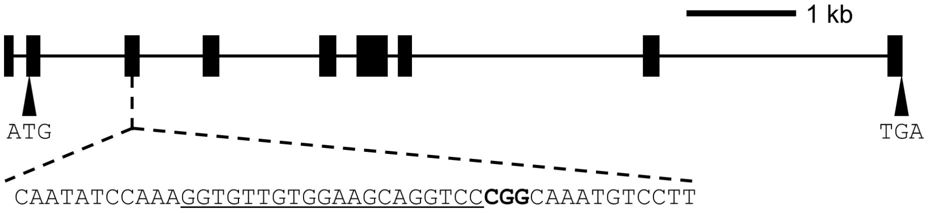


| | Lys | Gly | Val | Val | Glu | Ala | Gly | Pro | Gly | Lys | Cys | Pro | Phe | Glu | Thr |
|---|---|---|---|---|---|---|---|---|---|---|---|---|---|---|---|
| p50T | AAA | GGT | GTT | GTG | GAA | GCA | GGT | CC**C** | **GG**C | AAA | TGT | CCT | TTT | GAA | ACA |
| Δ5 | AAA | GGT | GTT | GTG | GAA | GCA | --- | --- | GGC | AAA | TGT | CCT | TTT | GAA | ACA |
| | | | | | A | | | | | | | | | | |
| In4 | AAA | GGT | GTT | GTG | GAA | GCA | GGT | CCC | GGC | AAA | TGT | CCT | TTT | GAA | ACA |
| | | | | | | | CCTG | | | | | | | | |

# D

*KWMTBOMO01920*

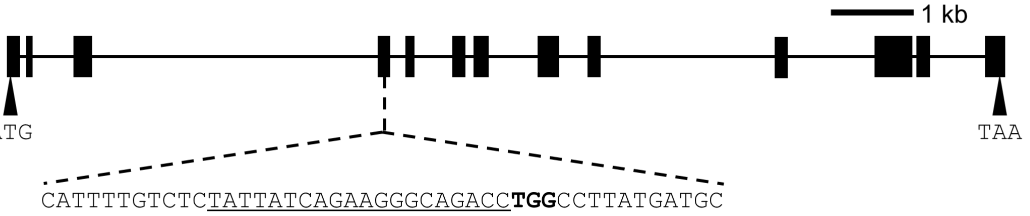


| | Leu | Tyr | Tyr | Gln | Lys | Gly | Arg | Pro | Gly | Leu | Met | Met | Arg | Ser | Leu |
|---|---|---|---|---|---|---|---|---|---|---|---|---|---|---|---|
| p50T | CTC | TAT | TAT | CAG | AAG | GGC | AGA | CC**T** | **GG**C | CTT | ATG | ATG | CGA | TCG | TTA |
| Δ2 | CTC | TAT | TAT | CAG | AAG | GGC | --A | CCT | GGC | CTT | ATG | ATG | CGA | TCG | TTA |
| Δ9 | CTC | TAT | TAT | CAG | AAG | GGC | --- | --- | --- | CTT | ATG | ATG | CGA | TCG | TTA |

# E

*KWMTBOMO01923*

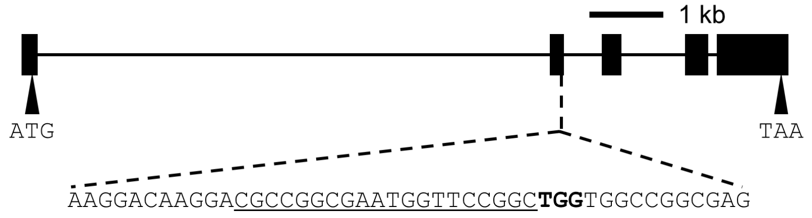


```
            Lys Asp Ala Gly Glu Trp Phe Arg Leu Val Ala Gly Glu Gly Asp
p50T        AAG GAC GCC GGC GAA TGG TTC CGG CTG GTG GCC GGC GAG GGC GAC

Δ6          AAG GAC GCC GGC GAA T-- --- -GG CTG GTG GCC GGC GAG GGC GAC

In11        AAG GAC GCC GGC GAA TGG TTC CGG CTG GTG GCC GGC GAG GGC GAC
                                        CATTCCACCAT
```

**Figure S2**: Mutations introduced to *l-t04* candidate genes by CRISPR/Cas9. The exon structure and gene sequences surrounding the guide RNA target site from wild types (p50T), *KWMTBOMO01913* (A), *KWMTBOMO01914* (B), *KWMTBOMO01917* (C), *KWMTBOMO01920* (D), and *KWMTBOMO01923* (E) KO homozygous mutants.

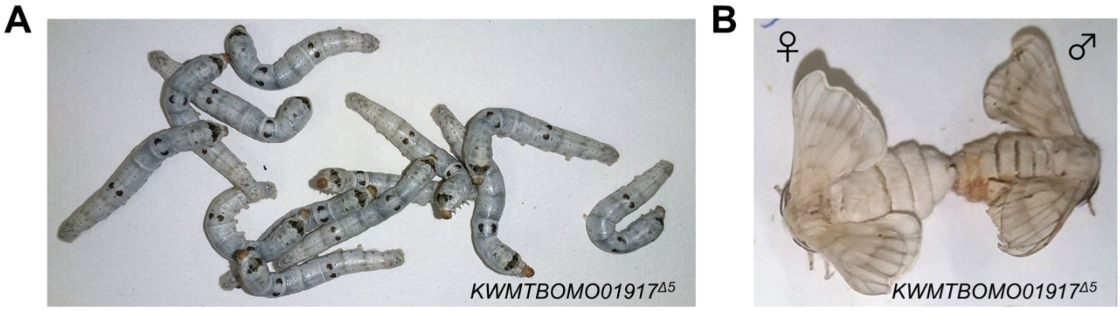


**Figure S3**: Fifth instar larvae (A) and moths (B) of *KWMTBOMO01917*$^{\Delta 5}$ KO mutants. *KWMTBOMO01917*$^{\Delta 5}$ mutants grew normally throughout their life stages.

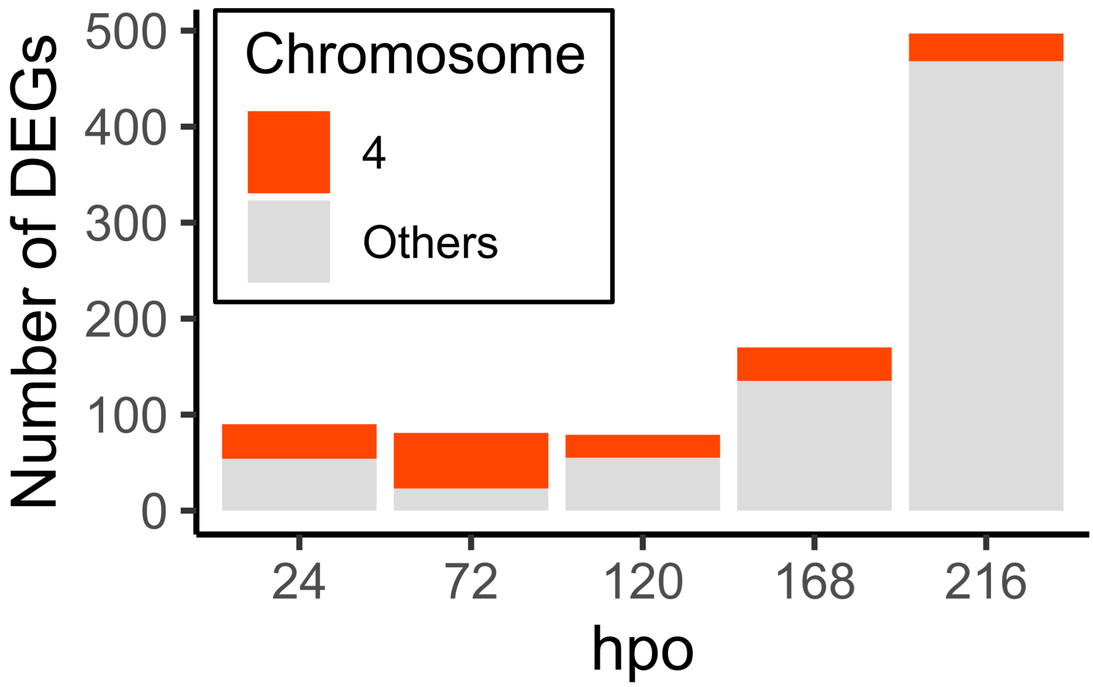


**Figure S4**: The number of differentially expressed genes (DEGs; FDR < 0.01) between wild type and *l-t04* mutant embryos. Several DEGs at 24, 72, and 120 hpo were located on chromosome 4, probably due to allele-specific expression differences between the *B. mori*- and *B. mandarina*-derived chromosomes. The number of DEGs located on other chromosomes increased from 168 hpo slightly before the lethal stage of the *l-t04* mutant (192–216 hpo).

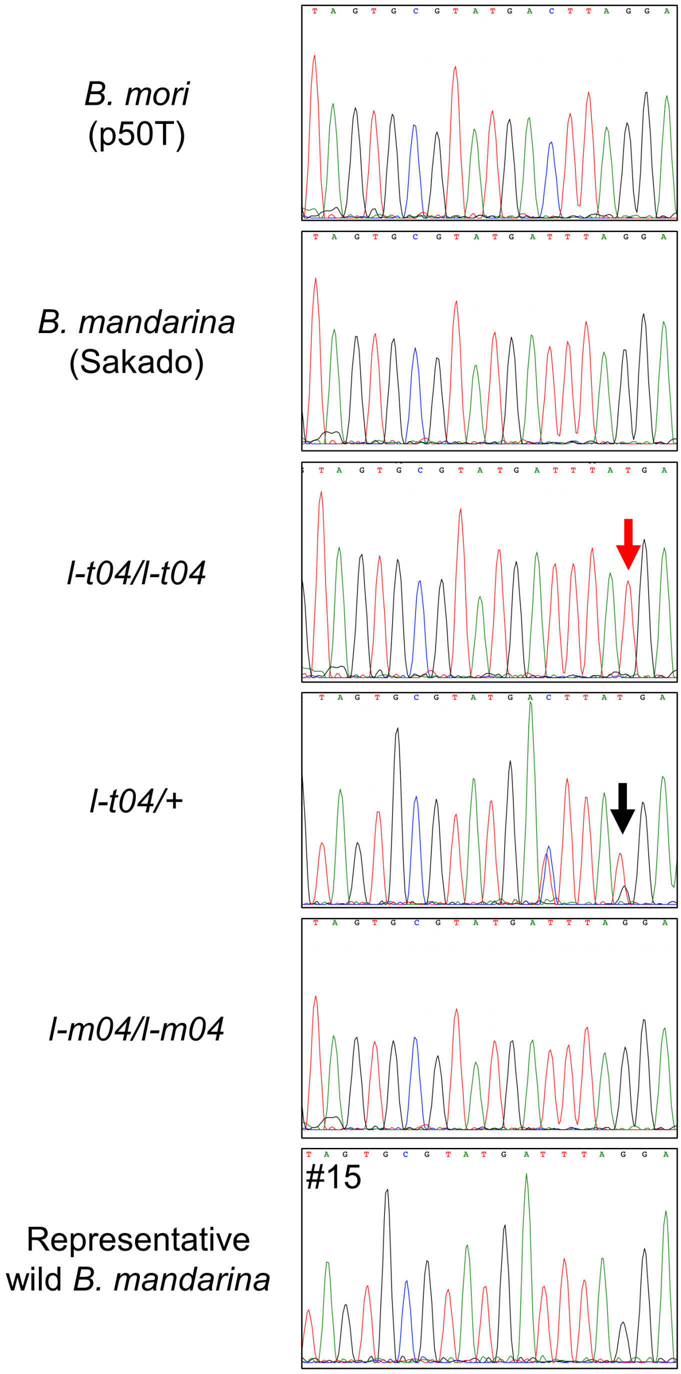


**Figure S5**: Sanger sequencing of a *Bmida* genomic region. A red arrow points to the homozygous G1969T mutation in *l-t04*/*l-t04,* and a black arrow indicates the heterozygous G1969T mutation in *l-t04*/+. The G1969T mutation is absent in parental strains *B. mori* p50T and *B. mandarina* Sakado and wild *B. mandarina* individuals.

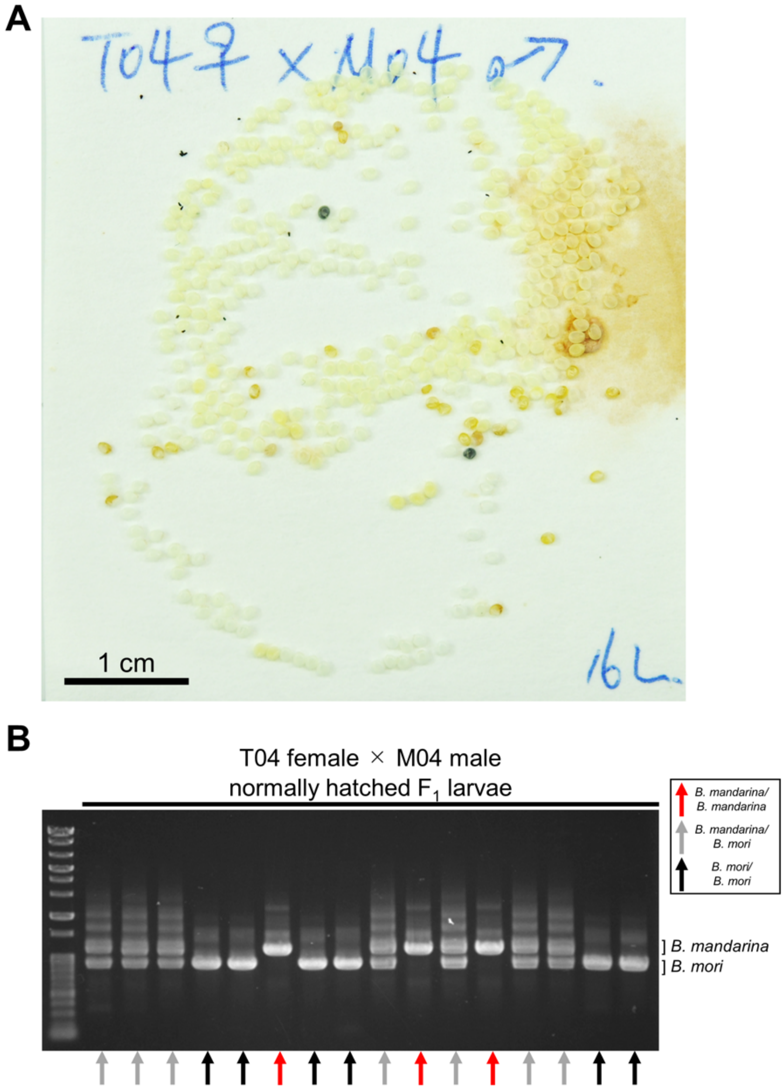


**Figure S6**: The *l-t04* and *l-m04* mutations are independent. (A) Post-hatch eggs from a crossing between T04 female and M04 male. (B) Genotypes of normally hatched T04 female × M04 male $F_1$ neonate larvae. Larvae homozygous for *B. mandarina*-derived chromosome 4 were not lethal. Genomic PCR was performed using a primer set closely linked to the *l-t04* responsible region (Bmo04-08m01 in Table S1). Individuals homozygous for *B. mandarina*, heterozygous, and homozygous for *B. mori* are indicated by red, gray, and black arrows, respectively. Gene Ladder Wide 2 (Nippon Gene) was used as a molecular size marker.

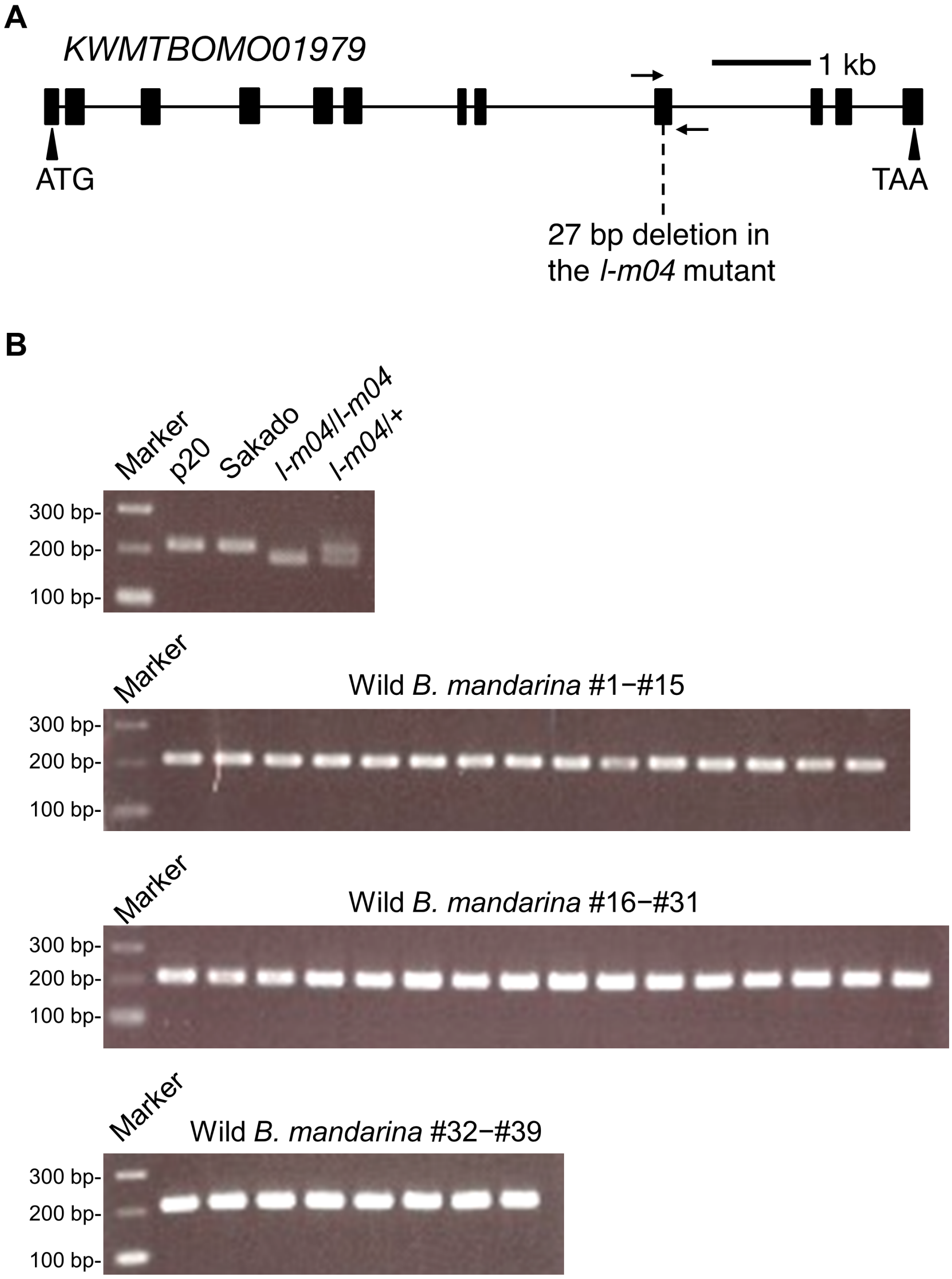


**Figure S7**: (A) The exon structure of *KWMTBOMO01979*. Arrows indicate the position of primers used in (B). (B) Gel electrophoresis of PCR products amplified from genomic DNA obtained from the wild-type *B. mori* (p20), *B. mandarina* (Sakado), *l-m04*/*l-m04*, *l-m04*/+ mutants (M04), and 39 wild *B. mandarina* individuals.

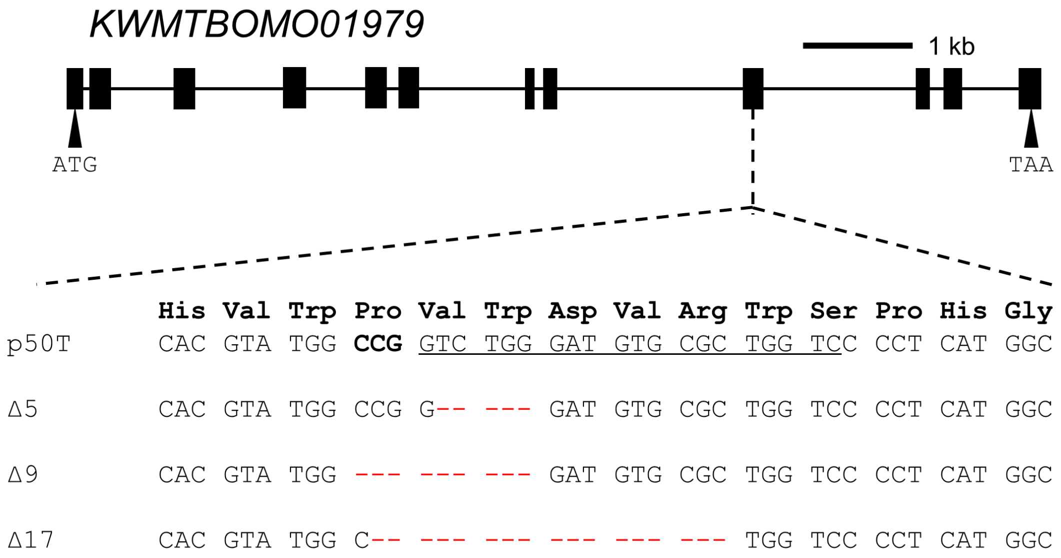


**Figure S8**: The exon structure of *KWMTBOMO01979* and *KWMTBOMO01979* gene sequences surrounding the guide RNA target site from wild-type (p20) and *KWMTBOMO01979* KO homozygous mutants.

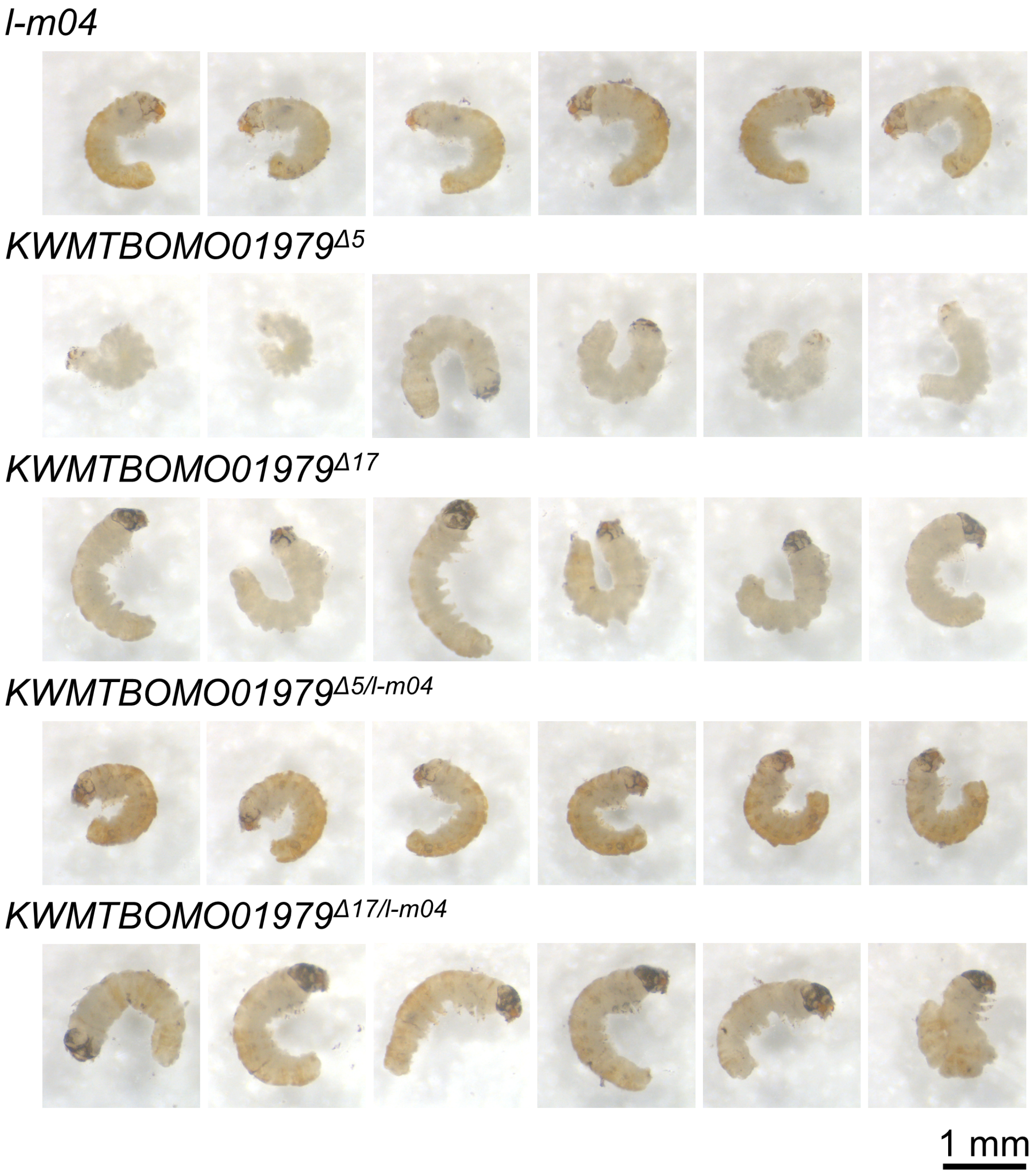


**Figure S9**: The lethal stages of the *l-m04*, *KWMTBOMO01979*$^{\Delta 5}$, *KWMTBOMO01979*$^{\Delta 17}$, *KWMTBOMO01979*$^{\Delta 5/l\text{-}m04}$, and *KWMTBOMO01979*$^{\Delta 17/l\text{-}m04}$ mutant embryos. The genotype of each embryo was confirmed by genomic PCR using the primers shown in Table S1. Scale bar: 1 mm.

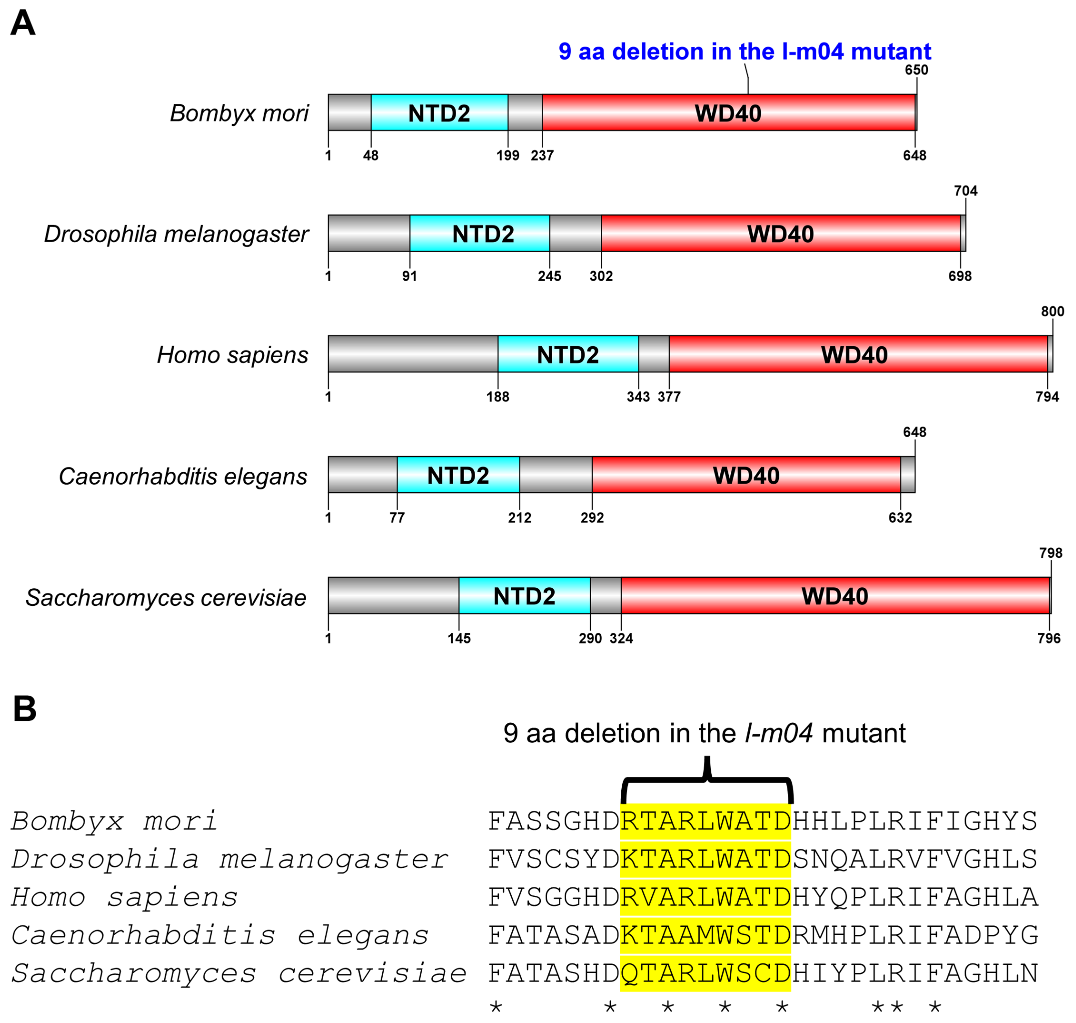


**Figure S10**: Protein structures and amino acid alignment of TAF5 homologs. (A) The structure of *B. mori* TAF5 (KWMTBOMO01979), *D. melanogaster* TAF5 (UniProt accession number P49846), *H. sapiens* TAF5 (UniProt accession number Q15542), *C. elegans* TAF-5 (UniProt accession number G5EF68), and *S. cerevisiae* TAF5 (UniProt accession number P38129). The protein domains were predicted using InterProScan (https://www.ebi.ac.uk/interpro/search/sequence/). NTD2: TFIID subunit TAF5, NTD2 domain. WD40: WD40/YVTN repeat-like-containing domain. (B) Alignment of TAF5 homologs containing the nine amino acid (aa) deletion regions in the *l-m04* mutant. The protein sequences were aligned using MUSCLE (Edgar, 2004).
